# Parasite histones mediate leak and coagulopathy in cerebral malaria

**DOI:** 10.1101/563551

**Authors:** Christopher A Moxon, Yasir Alhamdi, Janet Storm, Julien MH Toh, Joo Yeon Ko, George Murphy, Terrie E Taylor, Karl B Seydel, Sam Kampondeni, Michael Potchen, James S. O’Donnell, Niamh O’Regan, Guozheng Wang, Guillermo García-Cardeña, Malcolm Molyneux, Alister Craig, Simon T Abrams, Cheng-Hock Toh

## Abstract

Coagulopathy and leak, specific to the brain vasculature, are central pathogenetic components of cerebral malaria (CM). It is unclear how the parasite, *Plasmodium falciparum*, triggers these processes. Extracellular histones, released from damaged host cells, bind to cell membranes, causing coagulation activation, platelet aggregation and vascular leak in diverse critical illnesses. In CM patients, serum histones correlate with fibrin formation, thrombocytopenia, and endothelial activation; predicting brain swelling on MRI and fatal outcome. Post-mortem, histones bind to the luminal vascular surface, co-localizing with *P. falciparum*-infected erythrocytes (IE), and with thrombosis and leak. Purified *P. falciparum* histones or serum from patients with CM cause toxicity and barrier disruption in cultured human brain endothelial cells, reversed by anti-histone antibodies and non-anticoagulant heparin. These data implicate parasite histones as a key trigger of fatal brain swelling in CM. Neutralizing histones with agents such as non-anticoagulant heparin warrant exploration to prevent brain swelling and improve outcome.

## Introduction

Cerebral malaria (CM) is a severe complication of *Plasmodium falciparum* infection. Despite effective antimalarial drugs, 10-20% of children developing CM die (1), contributing to 400,000 malarial deaths per year, mostly in children in sub-Saharan Africa (2). Recent MRI studies implicate blood brain barrier (BBB) breakdown and brain swelling in the causal pathway to death (3, 4). Death typically occurs in the first 24 hours after admission (5), with children who do not reach critical levels of brain swelling frequently recovering rapidly. BBB stabilization, through targeting causal pathways to vascular leak in the brain, could halt this brain swelling and reduce mortality.

A defining feature of CM is cytoadherence of *P. falciparum* infected erythrocytes (IE) to endothelial cells (EC) and sequestration in the microvasculature (1)*. In vivo* retinal imaging (6, 7), post-mortem histology (8, 9) and *in vitro* data (10) demonstrate spatial-temporal links between sequestration and microvascular leak and thrombosis, and coagulopathy predicts fatal outcome in CM (11, 12). Post-mortem studies in African children demonstrate sequestration in multiple organs, whereas leak and coagulopathy are most prominent in the brain (9, 13, 14); implying that sequestration provides a parasite stimulus for vascular leak and coagulopathy and that the response to this stimulus is different in the brain (8, 15). The nature of this parasite stimulus remains unclear.

Extracellular histones, released by damaged or immune activated host cells have emerged as critical EC damage mediators in diverse severe illnesses including sepsis (16), inflammatory conditions (17) and trauma (18). Hallmark features of histone toxicity are thrombocytopenia (19) and microvascular thrombosis and leak (16, 18). In patients with sepsis or trauma, histone levels correlate with clinical severity scores (20), thrombocytopenia (19), coagulation activation (18, 20, 21) and predict outcome (22). In animal models of sepsis or trauma, the release of extracellular histones are causal in these processes and in fatal outcome, which are prevented by anti-histone antibodies (18, 20, 23), heparins, including non-anticoagulant heparins (24) (which neutralize histones) and by activated protein C (aPC, which degrades histones) (16). In mice, infusion of exogenous histones of >30mg/kg are toxic and of >60mg/kg are fatal; histologically histones are observed to bind to the endothelium, associated with microvascular coagulopathy and vascular leak (22). *In vitro*, histone binding to the EC membrane causes toxicity and barrier disruption (22, 23). The cationic domain of histones also induces Weibel Palade body exocytosis, endothelial activation and thrombocytopenia through platelet aggregation on von Willebrand Factor strings (25). Histones further induce a procoagulant phenotype through upregulation of endothelial tissue factor (26). By an unknown mechanism, histones decrease cell surface thrombomodulin *in vitro* (27), and induce thrombomodulin shedding *in vivo* (18).

Given the striking similarities between the vascular leak, coagulopathy and thrombocytopenia induced by histones in other conditions (16, 22) and those at sites of sequestration in CM, in particular the brain (8, 9, 14), we hypothesized that histones might be an important causal factor in CM pathogenesis. *P. falciparum*, as mammalian cells, contains histones (H2A, H2A.Z, H2B, H3, H4), packaged in nucleosomes with DNA. Following sequestration, intraerythrocytic merozoites multiply 16-24 times to form a schizont, increasing nuclear material, including histones, by an order of magnitude. Schizonts rupture releasing their contents, extruding *P. falciparum* histones *in vitro* into culture medium (28). Similar to mammalian histones, on cultured ECs, purified plasmodial histones cause inflammatory pathway activation, toxicity and barrier disruption (28). Therefore, histones may link sequestration and vascular pathology in CM; sequestration bringing histone-packed schizonts in contact with the endothelial surface, concentrating exposure to extruded histones many fold. The brain might be particularly vulnerable to this mechanism. Firstly, there are high levels of sequestration in the brain in CM (14, 29, 30). Secondly the brain may have reduced capacity to breakdown histones: the human brain has reduced innate capacity to produce activated protein C (aPC) (31), owing to low constitutive thrombomodulin and endothelial protein C receptor (EPCR) expression (32, 33), the receptors involved in aPC production. Moreover, parasite variants associated with the development of CM utilize EPCR as a binding receptor (34, 35), interfering with its function and the production of aPC (35, 36). Thus histones released by IE would be predicted to concentrate and be particularly toxic in the brain.

Supporting that *P. falciparum* histones may be released in patients with malaria, nucleosomes have been detected in the plasma of South-East Asian adults with malaria, which were higher in severe cases (28). However the association between nucleosomes (which have minimal toxicity (21)) and free histones is variable and it was not identified whether these nucleosomes were of host or parasite origin, or whether they were active. Thus, it remains uncertain whether significant levels of parasite histones are produced *in vivo* in patients with malaria and there are no data assessing the association between histones and clinical or laboratory indicators of severity or coagulation and leak, nor data to assess whether plasmodial histones bind in the vasculature at sites of sequestration.

Here we address these gaps. Using detailed laboratory, clinical and MRI imaging data we link histone levels in the blood to fibrin formation, endothelial activation and thrombocytopenia and to brain swelling and fatal outcome. Through post-mortem brain tissue samples from CM cases we show marked correlation between sequestration and the deposition of histones on the endothelial surface, and co-localisation with thrombosis and leak in the brain vasculature. We then demonstrate a causal role of *P. falciparum* histones in these processes through *ex vivo* experiments.

## Methods

### Patients and blood samples

Children aged 6 months – 16 years were recruited at Queen Elizabeth Central Hospital, Blantyre Malawi between January 2010 and August 2011. Inclusion criteria are described previously (8). Children who met WHO criteria for CM underwent funduscopic examination by an ophthalmologist: characteristic retinal changes indicate sequestration of IE in the brain(37) and distinguish retinopathy-positive CM with stringently defined CM (CM-pos) from cases with retinopathy negative CM (CM-neg), who are more likely to have an alternative diagnosis (1), to which malaria makes a variable contribution (38) and thus may have a different coma aetiology. Uncomplicated malaria and mild aparasitemic febrile illness cases were children with acute febrile illness without signs of organ compromise recruited from the hospital Accident and Emergency department (Emergency Room). Healthy controls were children attending elective surgery. Venous blood was collected at enrolment into plain or sodium citrate tubes and serum and plasma prepared as previously described (39), stored at −80°C until assays were performed. Circulating histone levels were quantified by a custom immunoblot assay (18–20) and Osteoprotegrin, Fibrin monomers, F1+2 fragment by ELISA as described previously (8, 11, 40).

There were no prior data on histone levels in CM on which to base a power calculation. The number of samples to be analysed was determined *a priori*, at the time of study design, based on the availability of samples and deemed to be appropriate based on comparison of histones levels in other conditions. All samples were processed and analysed together.

### MRI scans and scoring of brain swelling

MRI images were acquired using a 0.35-Tessla Signa Ovation Excite MRI scanner (General Electric). Images were scored independently by two radiologists who were blinded to patient disease group and outcome. A score from 1 – 8 was assigned to each scan, based on cerebral hemisphere swelling, using pre-specified criteria – described previously (3). We divided patients into 4 groups on the basis of this 8-point score: Score 1-3, No brain swelling; 4-5, mild brain swelling; 6, moderate brain swelling and 7-8, severe brain swelling. A number of children did not have MRI scans. When this was because they recovered from coma within 12 hours we deemed it likely that they did not have significant brain swelling and included them in category 1. Other MRI scans were not performed for several reasons (e.g. patient clinical unstable, equipment issues), we could not reasonably assign a category, and missing data were handled by listwise deletion.

### Isolation and purification of P. falciparum histones

ITG mature IE were lysed with saponin and *P. falciparum* histones (H2A, H2B, H3, H4) purified using a Kit (Active Motif). Protein concentrations were determined by Biorad Protein Assay, using bovine serum albumin and purified calf histones (Roche) standards and purity examined by SDS-PAGE and Coomassie staining (>95% pure; Fig. S3)

### Mass spectrometry sample preparation

Purified *P. falciparum* and human histones (New England Biolabs) (6µg), normal serum, histone spiked serum and CM patient serum were separated by 15% SDS-PAGE and stained with Coomassie brilliant blue. The excised gel slices (<35kDa) from SDS-PAGE, were cut into 1mm^3^ plugs, transferred to a microtube and fully de-stained using 25mM Ambic alternately with Ambic/MeCN (2:1). Cysteine reduction was performed by adding 100µL DTT solution (1.5mg/mL) and incubated at 60°C for 60 min. Samples were centrifuged and the supernatant was discarded. Alkylation was performed by the addition of 100µL iodoacetamide (10mg/mL) for 45 min (protected from light). Samples were centrifuged and the supernatant discarded. Gel plugs were then washed with Ambic (25mM) for 15min at 37°C. To fully dehydrate the gel plugs, samples were washed with MeCN. In-gel digestion was performed by adding 100µL of trypsin (12.5ng/µL in 25mM Ambic) to each sample with overnight incubation at 37°C, and reactions terminated by the addition of 10µL formic acid (1% final concentration). The solutions surrounding the gel plugs (containing the tryptic peptides) were retained for analysis. To extract additional peptides from the gel plugs, a further incubation with a solution containing water:MeCN:FA (50:49:1) and then MeCN:FA (80:19:1) was performed. Finally, solutions were pooled and dried to a 10µL solution.

### Liquid Chromatography-mass spectrometry analysis

Analysis was performed using an Ultimate 3000 RSLC™ nano system (Thermo Scientific, Hemel Hempstead), coupled to a QExactive-Hf™ mass spectrometer (Thermo Scientific). Samples were loaded onto a trapping column (Thermo Scientific, PepMap100, C18, 300 μm X 5 mm), using partial loop injection, for seven minutes at a flow rate of 9 μL/min with 0.1% (v/v) FA. Samples were then resolved on the analytical column (Easy-Spray C18 75 µm x 500 mm 2 µm column) using a gradient of 97% A (0.1% formic acid) 3% B (99.9% ACN 0.1% formic acid) to 60% A 40% B over 15 min at a flow rate of 300 nL min-1. The data-dependent program used for data acquisition consisted of a 70,000 resolution full-scan MS scan (AGC set to 1 x 10^6^ ions, with a maximum fill time of 20ms) the 10 most abundant peaks were selected for MS/MS using a 35,000 resolution scan (AGC set to 1 x 10^5^ ions with a maximum fill time of 100ms) with an ion-selection window of 3 m/z and a normalized collision energy of 28. To avoid repeated selection of peptides for MS/MS the program used a 15 second dynamic exclusion window. Sequence alignment was performed in PEAKs software (v8.5) against both *P. falciparum* and *Homo sapiens* databases. Once species-specific peptides were identified they were further verified using Skyline analysis software for quantification (comparisons between the specific amino acid sequences of *P. falciparum* and *Homo sapiens* histone proteins illustrated in Fig S4).

### Immunohistochemistry

Brain tissue samples of parietal cortex were collected at autopsy from Malawian children dying with encephalopathic illness and were formalin fixed and paraffin embedded as described previously (9). Based on clinical information and autopsy findings the cause of death was determined for each case by a clinical pathologist. We used samples classified into one of 3 overall categories as defined previously (1): 1) Definitive CM (CM1 and CM2) – children who met the case definition for CM during life and who at death had sequestration of IE in cerebral vessels and in whom no alternative cause of death was identified at autopsy; 2) ‘Faux CM’ (CM3) – met the case definition for CM during life but who had no visible sequestration of IE in cerebral vessels and in whom at autopsy another cause of coma and death was identified in all cases; 3) Aparasitemic non-malarial coma comatose patients who had no detectible malaria parasites in blood or tissue.

Cortical sections (4µm in thickness) were stained for histones and fibrinogen. Heat-induced antigen retrieval in citrate buffer (pH 6.0) was performed prior to incubation with primary antibodies: anti-histone H3 (Abcam); anti-Fibrinogen (Thermofisher)). Bound primary antibody was detected with an immunoperoxidase kit (EnVision Plus; Dako). Negative controls without primary antibody were used for all samples to confirm specificity. Immunohistochemistry was performed on all cases by a single investigator blinded to histologic diagnosis. Slides were scored by 3 investigators blinded to histological classification. 70 random vessels were scored from each slide. IE sequestration for each vessel was scored as: negative (0); positive but <50% of the vessel lumen (+) or >50% of the vessel lumen (++). Histone membrane staining for each vessel was scored as absent (0); weak (+) or strong (++). Fibrinogen extravasation as a marker of leak was scored for each vessel as absent or present.

There were no prior data on histone staining in post-mortem samples in CM or in other conditions. The numbers of samples to be stained and the numbers of vessels to be scored were based on numbers from previous studies comparing factors in CM. Numbers for staining and analysis were determined *a priori* and were not altered. Samples were stained together to avoid batch effects.

### Endothelial cell culture, endothelial cell damage assays and barrier function assays

Primary HBMEC (Cell Systems, US) were cultured in 1% gelatin-coated flasks, in Complete Medium containing 10% FBS (Cell Systems, US) as per manufacturer’s instructions.

For toxicity assays, HBMEC were treated with either purified histones in Cell Systems media with 2% serum or serum from healthy controls or patients (diluted 1:1 with PBS) for 1 hour at 37°C, under 5% CO_2_. Cell viability was determined by propidium iodide (PI) staining and quantified using flow cytometry. Cell toxicity in patient samples was calculated as the percentage of cells that were PI positive, subtracting the percentage of PI positive cells from the healthy donors from each sample. For anti-histone treatments, patient sera were pre-incubated for 10mins with anti-histone single-chain variable fragment (ahscFv; 200 μg/ml, synthesis described previously (18)) or with non-anticoagulant N-acetyl heparin (200 μg/ml; Sigma).

Transmembrane permeability of confluent HBMEC was analysed in a dual-chamber system (0.4 µM pore size; Millipore). HBMEC were treated with normal serum or patient serum (diluted 1:1 with PBS) for 1hr, replaced with horse radish peroxidase (HRP)-containing media. Leaked HRP over 1hr was determined using TMB substrate (ThermoFisher) on a microplate reader (450nm). Permeability was expressed as a fold change compared to monolayers treated with pooled normal serum from healthy UK donors [RETH000685].

Biological replicates were defined as independent experiments on primary HBMEC, treated with independently purified batches of purified histones or with serum from different patients. We generally used 3 biological replicates for experiments, based on routine practice for *in vitro* assays. For the 0 and 75ug/ml of purified histones and for the healthy control and CM patients with high histones we performed 6 and 7 biological replicates. Additional replicates were as controls for comparison with the anti-histone scFv, heparin and CM-low groups; the comparison with the other groups was similar to that presented and was significant after the initial 3 replicates.

### In vitro platelet aggregation

Platelets (2×10^3^/µl) prepared from healthy donors were mixed with pooled plasma spiked with malarial histones. Platelet aggregation was determined optically at 405nm (Multiskan Spectrum plate reader, ThermoScientific) in a 96-well plate, over 15mins at 37°C. To normalize for differences in optical density between plasma samples each sample was blanked with plasma in the absence of platelets, allowing the specific changes in optical density induced by platelet aggregation to be determined.

### Statistical analysis

Statistical analyses were performed using Stata (version 11; Statacorp) and Prism (version 8; GraphPad) software. Continuous variables were assumed to have normal or log normal distribution depending on their level of skewness. Differences between groups were compared using linear regression models. To adjust for multiple comparisons we used the Tukey (when comparing all groups to each other) or Dunnett tests (when comparing all groups to a control group). The association between histone levels and other variables was assessed by linear regression and expressed as correlation coefficients. For ordered categorical slide scoring data, the associations between histological classification, extent of sequestration and degree of fibrinogen extravasation were assessed by use of ordinal logistic regression models, controlling for clustering within cases and adjusting for any differences between scorers. All tests were two-tailed with a conventional 5% alpha-level.

## Results

### Circulating concentrations of extracellular histones are elevated in cerebral malaria cases and levels correlate with the degree of fibrin generation and with endothelial activation

Clinical characteristics of the patients are detailed in Table 1. Compared with CM-pos, CM-neg patients had a higher haemoglobin and platelet count and lower lactate level and parasite count. To explore whether histones are released *in vivo* and whether levels were associated with diagnosis, we measured circulating histones in serum samples taken from patients on admission. Histone concentrations were markedly higher in children with CM-pos than in children with CM-neg, non-CM encephalopathy, uncomplicated malaria, non-severe febrile illness or healthy controls (Fig 1A). These differences were not explained merely by an association with parasite density as there was only weak correlation between extracellular histone levels and peripheral parasite density (r=0.22 p=0.0044, Fig 1B) and there was no correlation between histone and histidine rich protein 2 levels (PfHRP2, a released parasite protein used as a marker of biomass [r=0.09, p=0.25]).

**Table 1.**
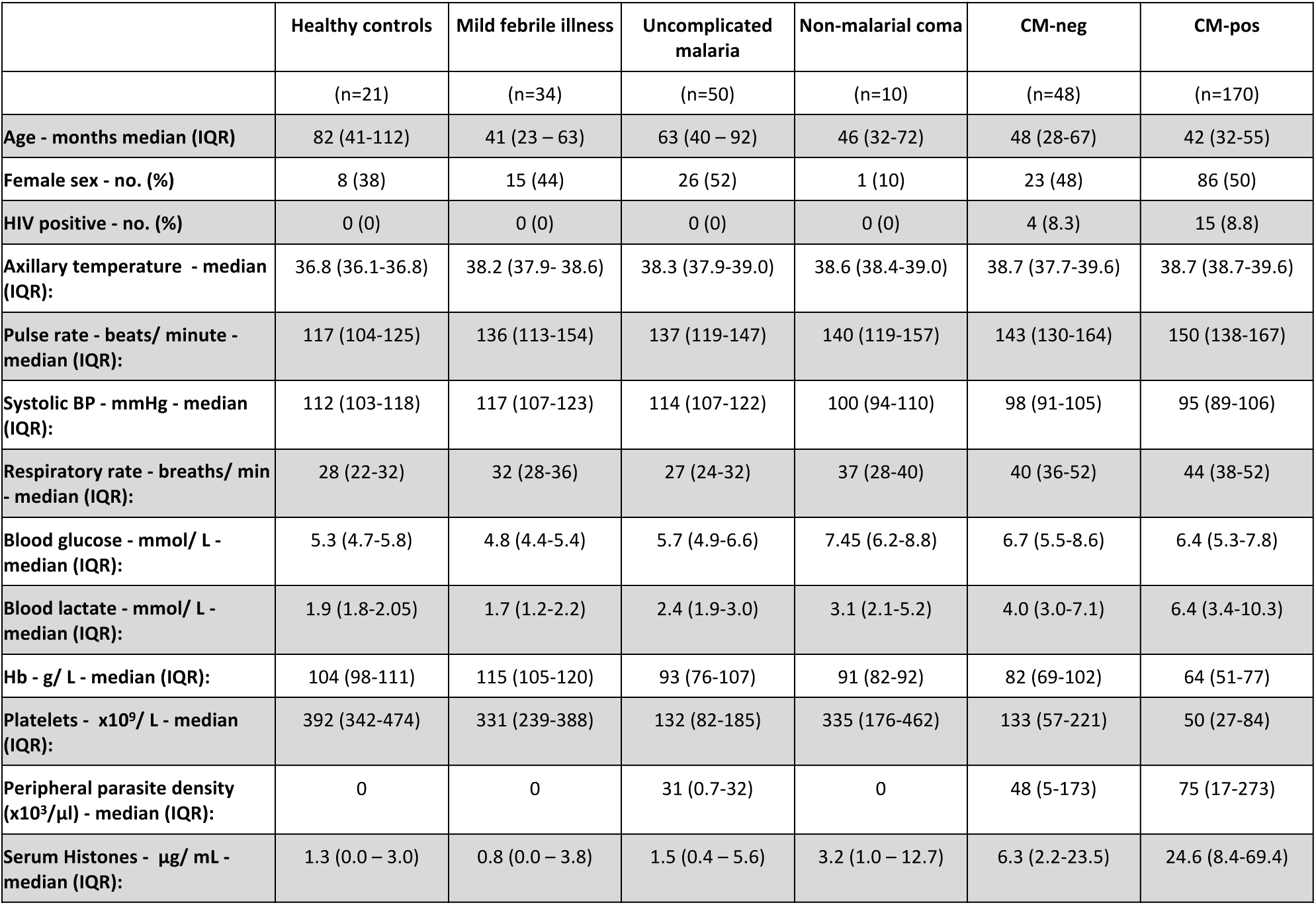
Clinical characteristics of the children. IQR - interquartile range; HIV - Human Immunodeficiency Virus; Hb - Hemoglobin.

|  | Healthy controls | Mild febrile illness | Uncomplicated malaria | Non-malarial coma | CM-neg | CM-pos |
| --- | --- | --- | --- | --- | --- | --- |
|  | (n=21) | (n=34) | (n=50) | (n=10) | (n=48) | (n=170) |
| Age - months median (IQR) | 82 (41-112) | 41 (23 – 63) | 63 (40 – 92) | 46 (32-72) | 48 (28-67) | 42 (32-55) |
| Female sex - no. (%) | 8 (38) | 15 (44) | 26 (52) | 1 (10) | 23 (48) | 86 (50) |
| HIV positive - no. (%) | 0 (0) | 0 (0) | 0 (0) | 0 (0) | 4 (8.3) | 15 (8.8) |
| Axillary temperature - median (IQR): | 36.8 (36.1-36.8) | 38.2 (37.9- 38.6) | 38.3 (37.9-39.0) | 38.6 (38.4-39.0) | 38.7 (37.7-39.6) | 38.7 (38.7-39.6) |
| Pulse rate - beats/ minute - median (IQR): | 117 (104-125) | 136 (113-154) | 137 (119-147) | 140 (119-157) | 143 (130-164) | 150 (138-167) |
| Systolic BP - mmHg - median (IQR): | 112 (103-118) | 117 (107-123) | 114 (107-122) | 100 (94-110) | 98 (91-105) | 95 (89-106) |
| Respiratory rate - breaths/ min - median (IQR): | 28 (22-32) | 32 (28-36) | 27 (24-32) | 37 (28-40) | 40 (36-52) | 44 (38-52) |
| Blood glucose - mmol/ L - median (IQR): | 5.3 (4.7-5.8) | 4.8 (4.4-5.4) | 5.7 (4.9-6.6) | 7.45 (6.2-8.8) | 6.7 (5.5-8.6) | 6.4 (5.3-7.8) |
| Blood lactate - mmol/ L - median (IQR): | 1.9 (1.8-2.05) | 1.7 (1.2-2.2) | 2.4 (1.9-3.0) | 3.1 (2.1-5.2) | 4.0 (3.0-7.1) | 6.4 (3.4-10.3) |
| Hb - g/ L - median (IQR): | 104 (98-111) | 115 (105-120) | 93 (76-107) | 91 (82-92) | 82 (69-102) | 64 (51-77) |
| Platelets - $\times 10^9$/ L - median (IQR): | 392 (342-474) | 331 (239-388) | 132 (82-185) | 335 (176-462) | 133 (57-221) | 50 (27-84) |
| Peripheral parasite density ($\times 10^3/\mu\text{l}$) - median (IQR): | 0 | 0 | 31 (0.7-32) | 0 | 48 (5-173) | 75 (17-273) |
| Serum Histones - $\mu\text{g}/\text{mL}$ - median (IQR): | 1.3 (0.0 – 3.0) | 0.8 (0.0 – 3.8) | 1.5 (0.4 – 5.6) | 3.2 (1.0 – 12.7) | 6.3 (2.2-23.5) | 24.6 (8.4-69.4) |

**Fig 1.**
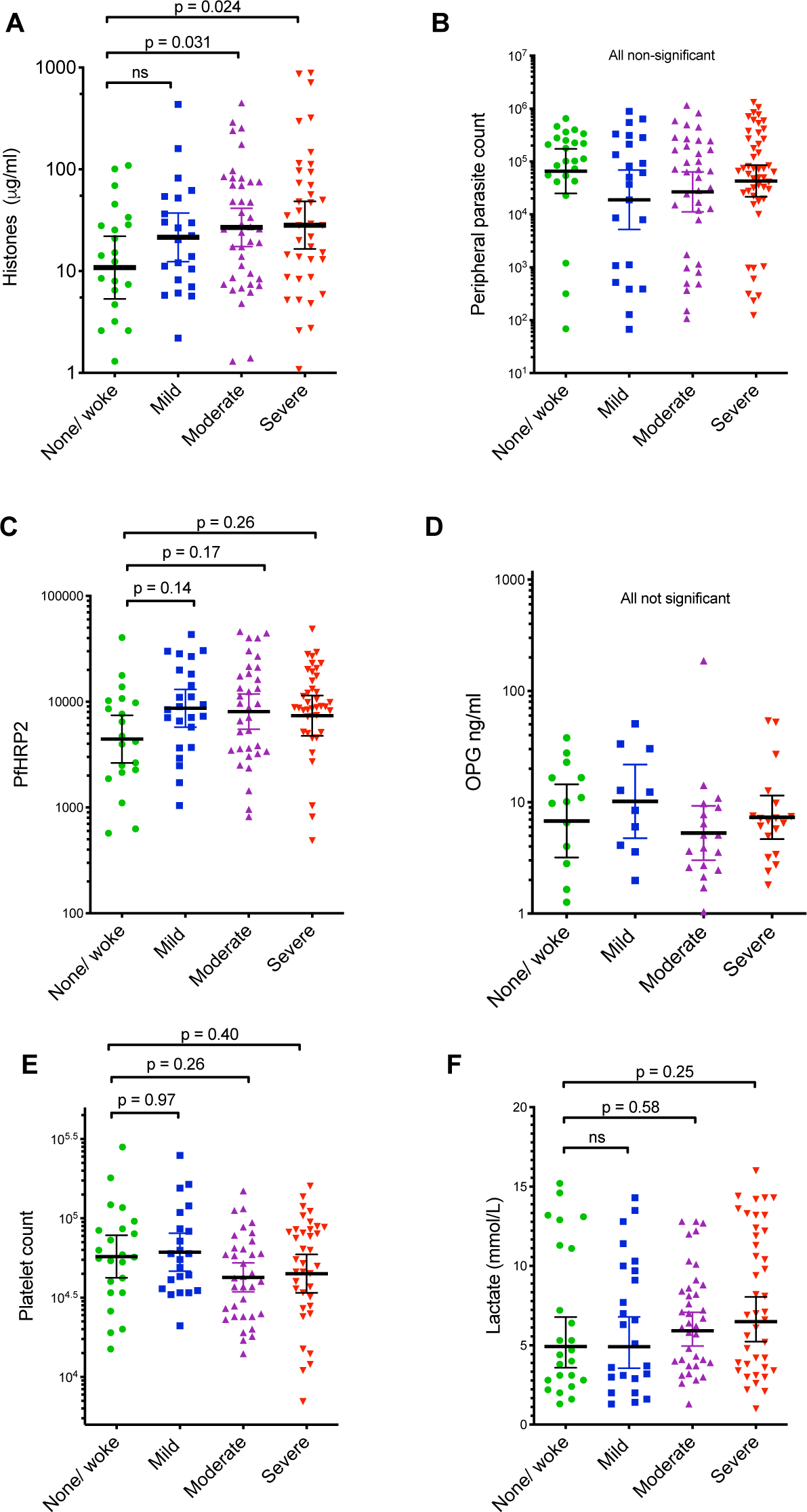
Circulating extracellular histones are elevated in cerebral malaria and correlate with intravascular fibrin generation and with endothelial activation. Extracellular histone levels were measured in serum samples taken on admission. (A) The mean concentration of extracellular histone levels in circulation was significantly higher in retinopathy positive cerebral malaria cases (CM-pos) than in all other patient groups including retinopathy negative CM (CM-neg). (B-F) correlations between serum extracellular histone concentration: peripheral parasite density in children with CM-pos (B); plasma fibrin monomer levels in children with CM-pos (C); prothrombin fragment F1+2 in children with CM-pos (D); platelet count among all children with CM (CM-pos and CM-neg) (E); plasma osteoprotegrin (OPG) concentration in children with CM-pos (F). HC = Healthy control; MF = Mild Febrile illness; UM = uncomplicated malaria; Non-CM (aparasitaemic children with encephalopathy [in coma] due to a cause other than malaria).

To explore histones as a possible trigger for coagulation activation in CM we assessed the association between circulating histones and markers of *in vivo* fibrin formation and coagulation activation (11). In CM-pos cases, plasma fibrin monomer concentrations correlated with circulating histone levels (r=0.56; p=<0.001, Fig 1C) more strongly than with (log) peripheral parasite density (r=0.34, p=<0.001), PfHRP2 (r=0.24, p=0.013), platelets (r=-0.18, p=0.2), lactate (r=0.33, p=<0.001), blood glucose (r=0.08 p= 0.58) or haemoglobin (r=0.06, p=0.89). Circulating histone levels showed a moderate correlation with prothrombin fragment F1+2 (a marker of thrombin generation (r=0.34, p=<0.001; Fig 1D)). Hence circulating histones better predict fibrin generation and coagulation activation than parasite density or other markers of disease severity.

Histones cause Weibel Palade Body (WBP) exocytosis and thrombocytopenia in mice through endothelial activation and increased platelet adhesion (25). Here circulating histone concentration correlated negatively with platelet levels, weakly in the subgroup of children with retinopathy positive CM (r= −0.22, p=0.0039 [in whom thrombocytopenia was nearly universal]), but moderately when patients with retinopathy negative CM were also considered (r= −0.41, p=<0.001; Fig 1E). Endothelial activation and WPB exocytosis are well established in CM including release of osteoprotegrin (OPG), which we have previously shown correlates with thrombocytopenia (40). Here circulating histone concentration correlated with plasma osteoprotegrin concentration (r = 0.54, p<0.001, Fig 1F). These data show a specific association between histones and CM-pos but not with CM-neg or aparasitaemic encephalopathy and suggest a link between extracellular histones and critical factors involved in clot formation and localization.

### Association between histone levels, brain swelling and fatal outcome

Given this association between histones and coagulopathy, a process implicated in brain swelling (41) and death (8, 42) in CM, we assessed the correlation between histone levels and fatal outcome and brain swelling. In children with CM-pos, the serum histone concentration was significantly higher in patients who died (n = 24; geometric mean 35.7 μg/ml [18.6–68.6 μg/ml]; Fig 2A) than in patients who survived (n = 146; geometric mean 21.6 μg/ml [16.4– 28.6 μg/ml]; p = 0.04).

**Fig 2.**
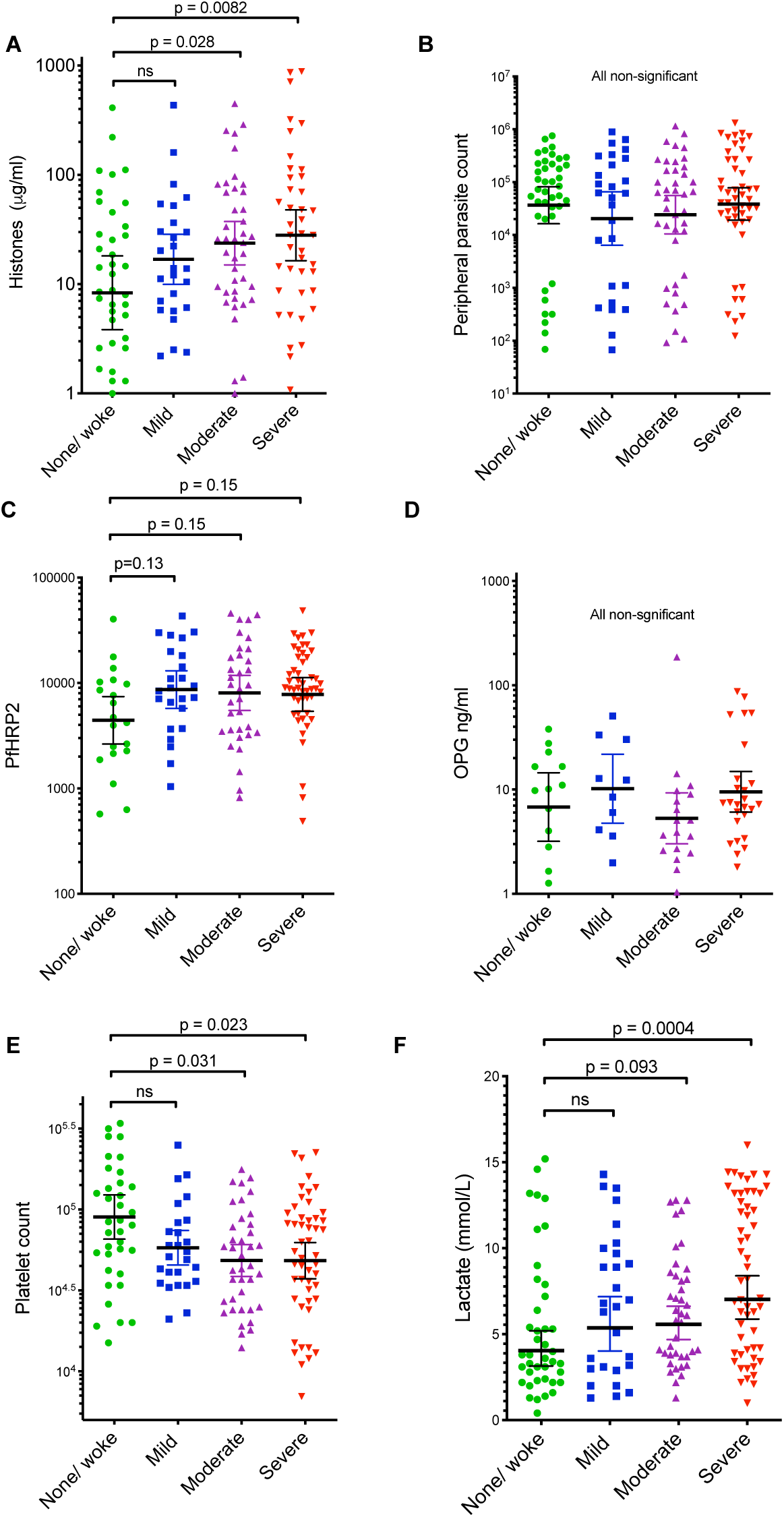
Extracellular histones are associated with fatal outcome and with the degree of brain swelling demonstrated on MRI scan. (A) In CM-pos cases (n=170), the mean extracellular histone level was higher in children who went on to die (fatal; n=24) than in those who survived (non-fatal; n=146). (B) Children were categorised by the degree of brain swelling on MRI; circulating histones were higher in children with moderate (n=41) or severe brain swelling (n=47) than in those with no evidence of brain swelling (n=22).

In CM-pos cases histone levels were 3 times higher in children who had moderate brain swelling (geometric mean 26.9 μg/ml; 95% CI 17.45 – 41.42, p= 0.031) or severe brain swelling (29.86 μg/ml; 95% CI 18.58 – 47.97, p = 0.024) than in children who had no evidence of brain swelling on MRI (8.79 μg/ml; 95% CI 3.09 – 25.01) (Fig 2B). In comparison peripheral parasite density, PfHRP2, lactate, platelet levels, and osteoprotegrin levels were not significantly associated with brain swelling (Fig.S1). There was a significant association between platelet levels and swelling and lactate levels and swelling when a less stringent definition of CM was used (i.e. when both CM-pos and CM-neg cases were included, Figure S2), this wider inclusion also increased the strength of association for histones (Figure S2). Taken together these data indicate a strong association between histone levels and the degree of brain swelling over and above other laboratory factors associated with severity in CM.

### Detection of significant levels of P. falciparum histones in patient samples using mass spectrometry

Owing to the highly conserved nature of histones, with >90% sequence homology between *Plasmodium* and human histones, available antibodies react with both human and *Plasmodium* histones (28). We developed a semi-quantitative mass spectrometry method (outlined in Figure 3A), to determine the proportion of parasitic and human histones within patient samples. Using *P. falciparum* histones purified from culture (Fig. S3), and pure human histones, we identified specific peptides for both H4 (Fig. 3B, C, Fig. S4) and H2A.Z (Fig. S4) that distinguished between *P. falciparum* and human histones (Fig. 3D, E). We then applied this method to serum samples from 10 children with CM-pos. *P. falciparum* and human histones were identified in all 10 CM cases, with *P. falciparum* histones constituting a mean of 51% (range 2% to 91%, Fig. 3F, G) of the total histone concentration.

**Figure 3.**
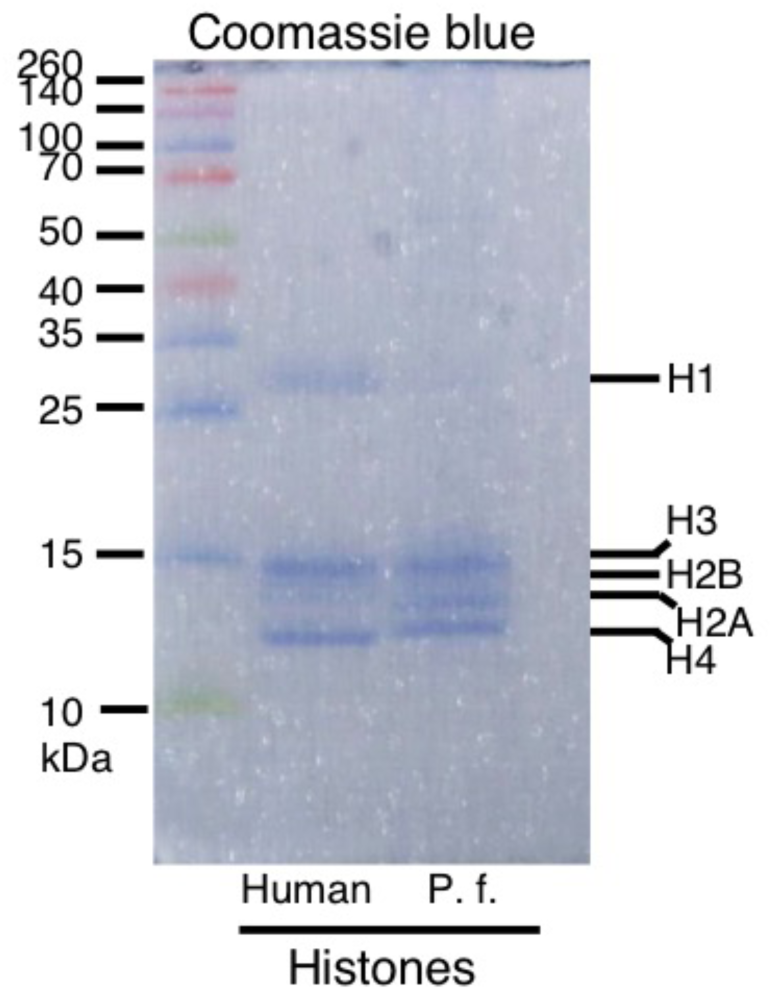
Mass spectrometry analysis of origin of extracellular histones in cerebral malaria cases. (A) Schematic representation of the methodology used for isolation, purification and mass spectrometry analysis. (B,C) Using Skyline software and by aligning trypsin fragments to reference amino acid sequences we were able to identify specific histone H2A.Z and H4 peptides that were present in purified *P. falciparum* (malarial) preparations, that were not present in purified human histones (H1, H2A, H2B, H3 and H4) and vice versa. Typical peptides are presented from human (B) and malarial (C) database searches. Using Skyline software, we were able to identify histone H4 peptides for each species that demonstrated different Mass/Charge ratios with distinct human and *P. falciparum* peptides and also distinct H2A.Z human and *P. falciparum* peptides (data not shown). (D,E) This enabled us to identify with high specificity and *P. falciparum* (D) and Human (E) species-specific peptides derived from samples spiked into PBS (left) or serum (right); data shown are for H4. (F) In CM-pos patient serum (n=10) we were able to *P. falciparum* histones H2A.Z and H4 in the samples as well as human H2A.Z and H4 (data not shown). (G) We combined the contribution of these two components to estimate the variable proportions of circulating human and *P. falciparum* in the patient serum, demonstrating a significant contribution of *P. falciparum* histones to the total pool.

### Accumulation of histones at the endothelial surface in the brain in fatal cases is associated with sequestration and with blood brain barrier breakdown

Histone mediated barrier disruption is caused by histones binding to the endothelium, observed by histology in histone-infused mice (22). To explore whether extracellular histones bind to the endothelium in CM we performed immunostaining for histones in post-mortem brain samples from Malawian children (details of cases in Table S1). Compared with “faux CM” (CM3) cases (n=6, Fig 4A) or non-CM cases (n=5) luminal histone staining was more frequent and stronger in CM cases (CM1/2, n=15, Fig 4B). Quantifying this by scoring with observers blinded to diagnosis, strong membrane staining was markedly associated with definitive CM when compared with faux CM (odds ratio [OR] 2.6; 95% Confidence Interval [CI] 1.7 – 3.9; p<0.001) or non-CM (OR 7.2; 95% CI 5.0 – 10.6; p<0.001; Fig 4C).

**Figure 4.**
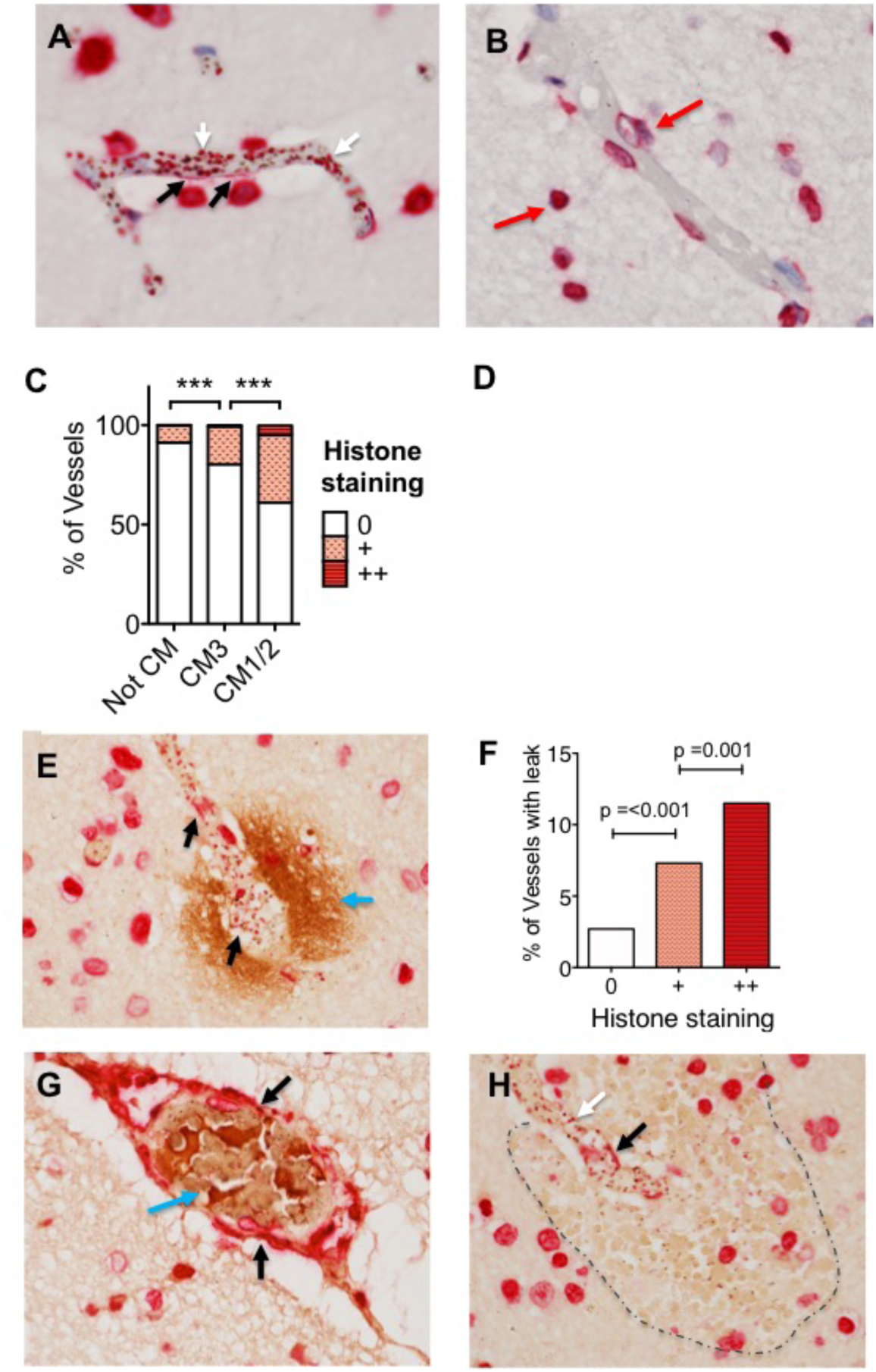
Histones accumulate at the endothelial surface in the cerebral microvasculature, associated with sequestration, coagulopathy and blood brain barrier breakdown. (A) Cerebral malaria case showing histone staining in close proximity with endothelial cell luminal surface (black arrows) and in both mammalian nuclei and malaria infected red blood cell (IE) nuclei (white arrow); (B) Non-CM case with no histone endothelial membrane binding, histone staining can be seen in mammalian cell nuclei (red arrows); (C) Extracellular histone staining is markedly increased in CM1/2 “true cerebral malaria”; (n=15) compared to ‘faux CM’ cases (peripheral parasitaemia, no sequestration in the brain and another cause of death at autopsy, CM3; n=6) or aparasitaemic non-CM cases (n=5); (D) In CM cases (CM1/CM2; n=15) there is a strong association between the degree of sequestration and the presence and strength of histone membrane staining. (E) Histone endothelial membrane staining (black arrows) co-localizing with fibrinogen extravasation (blue arrow), which is indicative of blood brain barrier breakdown. (F) Strong association between the extent of histone endothelial membrane staining and the presence of fibrinogen extravasation. (G) Histone membrane staining (black arrows) co-localizing with thrombosis (blue arrow). (H) Histone membrane staining (black arrow) co-localizing with a ring haemorrhage (edge demarcated by dotted line).

Among definitive CM cases there was a strong association between histone membrane staining and the presence of IE. This increased with more intense IE-sequestration: when sequestration was present but in less than 50% of the vessel (+) the OR of histone membrane staining being present was 5.2 (95% CI 2.8 – 9.7, p<0.001; Fig 4D); when greater than 50% of the vessel contained sequestered IE (++) the OR for the presence of histone staining was 16.9 (95% CI 9.2 – 31.3; p<0.001).

Histone staining was also strongly correlated with areas of BBB breakdown, demonstrated by staining for fibrinogen extravasation (Fig 4E): weak histone staining was associated with an OR of 2.8 for the presence of fibrinogen extravasation (95% CI 1.6 – 5.0; p=<0.001, 4F) and strong histone staining with an OR of 4.5 for fibrinogen extravasation (95% CI 1.8 – 11.4; p=0.001), as shown in fig.4H as “% of vessels with leak”. Histone staining was also observed to co-localize with thrombi (Fig 4G) and with ring hemorrhages (Fig 4H).

### Purified P. falciparum histones and serum from CM cases induce endothelial damage and barrier disruption

Mammalian histones directly induce endothelial cell membrane damage and barrier disruption on human vein umbilical vein EC (16, 18) and *P. falciparum* histones induce damage in dermal and lung EC (28). To investigate the potential relevance of this in the brain, we tested whether purified *P. falciparum* histones cause cell damage and leak on primary human brain microvascular EC (HBMEC). *P. falciparum* histones induced significant cellular toxicity (Fig. 5A, B) similar to the effects seen with mammalian histones (16, 18). To demonstrate that this effect was specifically induced by histones, and not a contaminant, we used an anti-histone single-chain Fragment variable (ahscFv), previously shown to inhibit histone toxicity (18, 43). ahscFV abrogated histone-induced toxicity (Fig. 5A). Non-anti-coagulant heparin, a potential treatment with minimal toxicity that prevents toxicity of mammalian histones (24), also prevented *P. falciparum* histone toxicity on HBMEC (Fig. 5A).

**Figure 5.**
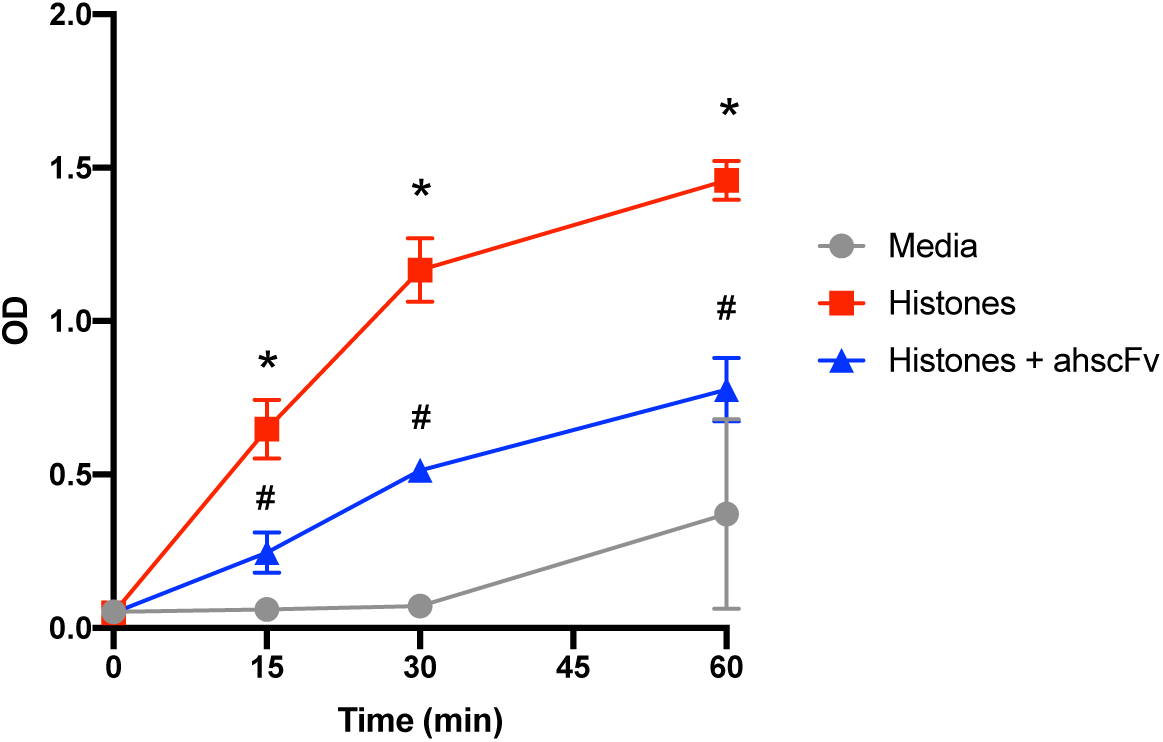
*P. falciparum* histones induce endothelial cell damage, permeability and platelet aggregation. A) HBMECs were treated for 1 hour with medium with or without purified *P. falciparum* histones (conc) ± anti-histone single-chain Fragment variable (scFv) or non-anticoagulant heparin. Cell toxicity was determined by propidium iodide staining using flow cytometry. Data are expressed relative to cells treated with media alone (set to 0%). ANOVA test * = p < 0.05 when compared with untreated, # = p < 0.05 when compared with CM-pos high treated with histone alone; B) HBMECs were treated for 1 hour with serum from retinopathy positive CM (± anti-histone scFv), uncomplicated malaria, mild non-malarial febrile illness, non-malarial encephalopathy or retinopathy negative malaria and healthy controls. Cell toxicity (means ± SD) relative to HBMEC treated with serum from healthy control cases (set to 0%) are presented. C) Transwell permeability changes of HBMEC monolayer are expressed as fold changes in HRP pass through compared to cells treated with normal healthy serum. *ANOVA test shows a significant decrease compared with normal (P < 0.05), #p < 0.05 when compared with retinopathy positive CM alone D) Platelet rich plasma was incubated with different concentrations of *P. falciparum* histones ± anti-histone scFv. Platelet aggregation (%) (means ± SD;) are presented following 15 mins incubation. ANOVA test, *p < 0.05 when compared with untreated, #p < 0.05 when compared with that treated with histone alone. 3 biological replicates were used in all experiments except for 0ug/ml and 75ug/ml in A) (6 replicates; additional replicates as controls for heparin and anti-histone scFv assays and for CM-pos high and HC in B) (7 replicates; additional replicates performed for comparison with scFv and CM-pos with low histones).

To investigate whether circulating histones from patients induce membrane toxicity, we incubated patient serum with HBMEC. Serum from CM-pos cases with elevated histones (histone concentration >100ug/ml; Fig 5B; n=5) induced significant cellular toxicity, whereas serum from CM-pos cases without substantially elevated histones levels (histone concentration <25ug/ml; n=3) did not, nor did samples from children with uncomplicated malaria (n=3), mild non-malarial febrile illness (n=3), non-malarial encephalopathy (n=3) or retinopathy negative CM (n=3; Figure 5B). Serum-induced toxicity was abrogated by ahscFv treatment, supporting a causal link with histones in the serum (Fig. 5A, B).

We next investigated the effect of purified *P. falciparum* histones and patient serum on barrier integrity. Similar to human histones, *P. falciparum* histones induced rapid barrier disruption in HBMEC. This leak was reversed by ahscFv (Fig S5). Similarly, serum from CM-pos cases with high histone levels (n=3) induced leak, but serum from CM-neg cases and other control groups (all n=3) did not. Leak in the CM-pos cases was abrogated by ahscFV, also supporting that histones in the serum were causal in this leak (Fig. 5C).

Given the correlation between histones and thrombocytopenia in CM (Fig. 1E) we investigated whether *P. falciparum* histones also cause platelet aggregation. Incubation of purified *P. falciparum* histones with platelet rich plasma from normal healthy controls resulted in dose dependent platelet aggregation, inhibited with ahscFv treatment (Fig. 5D).

## Discussion

A number of factors released from IE have been shown to cause endothelial damage or leak *in vitro* including glycosylphosphatidylinositol (44), extracellular vesicles (45), hemozoin and PfHRP2. IE-EC receptor-ligand interactions also cause endothelial perturbation (46–48). While it seems likely that CM pathogenesis constitutes a combination of interacting factors, rather than a single toxin or ligand (49, 50), we sought a factor that is necessary for CM vascular pathology and targetable with a safe and deployable treatment. Histones were a compelling candidate. Firstly because of the strong parallels between the clinicopathological features of histone-induced vascular pathology in other conditions and those in malaria. Secondly, because the sequestration of histone-packed IE in tissues would predict substantial concentration of histones being extruded to the endothelial surface. Thirdly because histones are a plausible target for an adjunctive therapy; treatments targeted against histones are protective in animal models of sepsis and trauma, even though extracellular histones are clearly not the sole factor contributing to pathogenesis in either of these conditions.

Our data provide evidence for histones as a mediator of the vascular pathology in the brain in CM that link causal data from *ex vivo* experiments (patient serum directly causes leak and toxicity, which is reversed by blocking histones) to multi-model observations in a rigorously defined patient cohort. Correlation between histones levels, diagnosis, fatal outcome, thrombocytopenia and fibrin production imply a role for histones in death and in key pathogenetic processes. Employing MRI scans and a grading system we established a correlation between serum histone levels and the level of brain swelling in children. We then showed that a significant proportion of histones were of parasite origin. Although human histones are also toxic, they are produced by diverse activated or damaged cells and might be a bystander event, triggered distant from sites of sequestration and vascular pathology. In contrast, parasite histones in the systemic circulation strongly suggest downstream detection of histones released from rupturing mature schizonts, in which histones are concentrated 16– 24-fold, occurring almost exclusively in sequestered IE. Examination of histological staining in post-mortem CM brain samples supported this paradigm. Extracellular histones were bound to the EC membrane, more frequently in CM cases than controls and spatially associated with the presence of sequestered IE and with areas of fibrinogen leak and thrombosis.

To confirm whether plasmodial histones might be causal in these pathogenetic events, we purified *P. falciparum* histones from parasites grown in culture and showed that they induced membrane damage and leak in primary HBMEC and platelet aggregation in platelets from healthy donors. These effects were prevented by specific ahscFv. Patient serum from CM cases with high levels of histones also induced EC membrane toxicity and leak. Both were blocked by pre-incubation with ahscFv, indicating that the effects were caused by active histones in serum. Heparins, including non-anticoagulant heparins have been shown to neutralize the effects of mammalian histones and may represent promising therapies. As a proof of concept, we showed that non-anticoagulant heparin prevented toxicity from *P. falciparum* histones. Taken together, these data show that *P. falciparum* histones are produced at significant levels *in vivo*, that they circulate in an active form, show a causal role for histones from patient serum samples *ex vivo* in processes leading to CM pathogenesis and provide multiple points of evidence supporting a role of histones in key disease processes in patients.

The locations of plasmodial histone production and what we know about modifiers of histone response fit well with the non-uniform pattern of vascular involvement in CM, whereby coagulopathy and leak are localized to sites of IE sequestration and in particular to the brain. It is notable that the median concentration of histones in the serum in CM-pos cases was 24.6μg/ml, and that toxicity to HBMEC in our assay was only seen at histones concentrations of >50μg/ml (similar to mammalian histones and to experiments using purified exogenous histone infusion in mice (16, 22, 28)). The implication being that in most patients with CM, histone levels in the circulation do not reach levels sufficient to cause systemic toxicity. This is in keeping with the observed clinical pattern of disease in CM in African children: deep coma and marked cerebral irritability, generally without multi-organ failure (14) or systemic coagulopathy (11). In contrast it seems highly plausible that *P. falciparum* histones concentrate several-fold at sites of intense sequestration (Fig. 3B) and cross this toxic threshold. We hypothesize that the brain is particularly vulnerable to histone toxicity because of reduced capacity to produce aPC. This would not be expected to manifest in conditions involving release of histones from immune-activated cells such as in sepsis and trauma, given that the brain is an immune-privileged site (51). The paradigm in CM is different; parasite histones reach high levels in the brain through IE sequestration. Moreover, IE sequestration in the brain may itself impair aPC production - firstly because IE reduce surface thrombomodulin and EPCR, putatively by receptor cleavage (8, 15). Secondly, parasite variants associated with the development of CM (expressing domain cassette 8 [DC8]) reduce aPC production, by binding to EPCR and inhibiting its activity (52). DC8 variants also show a tropism for brain endothelium (34, 52, 53). Hence parasites in CM patients may be more likely to concentrate plasmodial histones in the brain, through sequestration, and simultaneously may prevent their breakdown, through inhibiting aPC production. In support of this, DC8 expressing variants are associated with both thrombocytopenia and brain swelling (48); aPC inhibition potentially increasing both histone-induced platelet aggregation and histone-induced endothelial leak. It is notable that histones are implicated in neurotoxicity and ischemic damage in neurodegenerative conditions and stroke, and that in animal models these effects are reversed by aPC (54–56).

Our study has several limitations. Firstly, our study is in human patients. While generally a strength, this leads to marked heterogeneity, including in variables that might affect histone levels, such as length of illness and timing of antimalarial administration. Further we took blood from each patient at only one timepoint, representing a snapshot in a dynamic disease process. This precluded examination of the temporal association between histone levels and other variables. Secondly, while the association between histone binding and sequestration and the finding that 51% of histones in serum were of parasite origin are both highly suggestive of a parasite origin for luminal histones, we did not prove this. Nonetheless concentration of host histones at sites of IE sequestration would also be predicted to have similar effects and to respond to similar treatments.

Given that a significant proportion of histones detected in blood are of parasite origin it is notable that histone levels do not correlate well with parasitemia or PfHRP2. This may reflect the limitations of each of these assays, used at a single time point, to determine total parasite biomass. Firstly, our main assay to determine histone levels does not distinguish human from parasite histones. Serum histone levels are likely to be a function of production, breakdown and luminal binding and hence it is unclear how accurately serum histone levels of either species correlate with total production. Secondly peripheral parasitemia is a poor predictor of total parasite biomass: sequestered IE do not circulate, and so the concentration of parasites detectable in the periphery fluctuates markedly depending on the stage of the majority of the parasites in an individual patient. Thirdly, PfHRP2, a soluble parasite factor, has a long half-life and therefore its concentration in serum is a function of parasite biomass and duration of infection. While PfHRP2 is a predictor of parasite biomass and correlates with disease severity in several populations (57, 58), among Malawian children with CM, serum PfHRP2 levels do not correlate well with markers of severity (such as lactate or thrombocytopenia) or with outcome (59).

Further work is warranted to explore the biology and timing of plasmodial histone release and the mechanism of action of plasmodial histones in greater detail. A specific antibody against *P. falciparum* histone would be useful to differentiate *P. falciparum* histone levels in serum and in tissue. It remains to be determined whether agents that neutralize or degrade histones can reduce brain swelling during the critical 24 hours after hospital admission and thereby improve outcome in CM. Potential agents include aPC or heparin (24, 60, 61). Modified non-anticoagulant heparins are a rational first choice, particularly given their use in critically ill patients with a variety of inflammatory diseases (61) and in patients with sickle cell crisis (62). There is a planned phase II study in patients to use a modified heparin to reverse binding and rosetting in malaria. A different dosing regimen is likely to be needed to reverse the effects of histones than to block binding, which would require further investigation. However, the possibility that modified heparins could be synergistic in malaria – both reducing binding and neutralizing heparins – make the potential benefits more compelling. Finally, since cells in all eukaryotic organisms contain histones it will be important to explore whether parasite histones contribute to pathogenesis in other parasitic infections.

## Acknowledgments

For recruiting and caring for patients, we would like to thank the nurses and clinicians on the Paediatric Research Ward team (Malawi–Liverpool–Wellcome Clinical Research Programme and Blantyre Malaria Project), and the nurses and clinicians in the Department of Paediatrics and Child Health (Queen Elizabeth Hospital, Blantyre, Malawi). For performing the fibrin-based assays, we would like to thank C. Powell (Roald Dahl Haemostasis & Thrombosis Centre, Liverpool, U.K.). For histopathology advice we would like to thank Dan Milner. For technical support with histopathology we would like to thank Qian Zhen (Program in Dermatopathology, Department of Pathology, Brigham and Women’s Hospital, Boston, U.S.A.). For providing laboratory space and advice we would like to thank Dyann Wirth. For helpful comments on the manuscript we would like to thank Andy Waters and Matthias Marti (Wellcome Centre for Integrative Parasitology, University of Glasgow).

## Funding

This work was supported by funding from the Wellcome Trust (to C.A.M; 109698/Z/15/Z) and Academy of Medical Sciences (to C.A.M.); and a grant from the NIH (T. E. Taylor, 5R01AI034969-14). The Malawi–Liverpool Wellcome Clinical Research Programme is supported by core funding from The Wellcome Trust (084679/Z/08/Z).

## Author contributions

C.A.M., S.T.A., A.G.C., W.G. and C.H.T., conceived the study and designed experiments. C.A.M. and S.A. performed analysis. Y.A., S.A., J-Y. K., J.S., J.M.T., N.O. and C.A.M. performed laboratory experiments. K.B.S. and T.E.T. ran the clinical study and provided clinical and scientific input. M.E.M. provided clinical and scientific input. G.M., G.G-C. and J.O. designed experiments and provided scientific and technical input. C.A.M. wrote the original draft with significant input from A.G.C, S.A. and C.H.T. All authors contributed to critical review and editing of the manuscript.

## Competing interests

The authors have no conflicting interests.

## Supplementary Material

**Fig S1.**
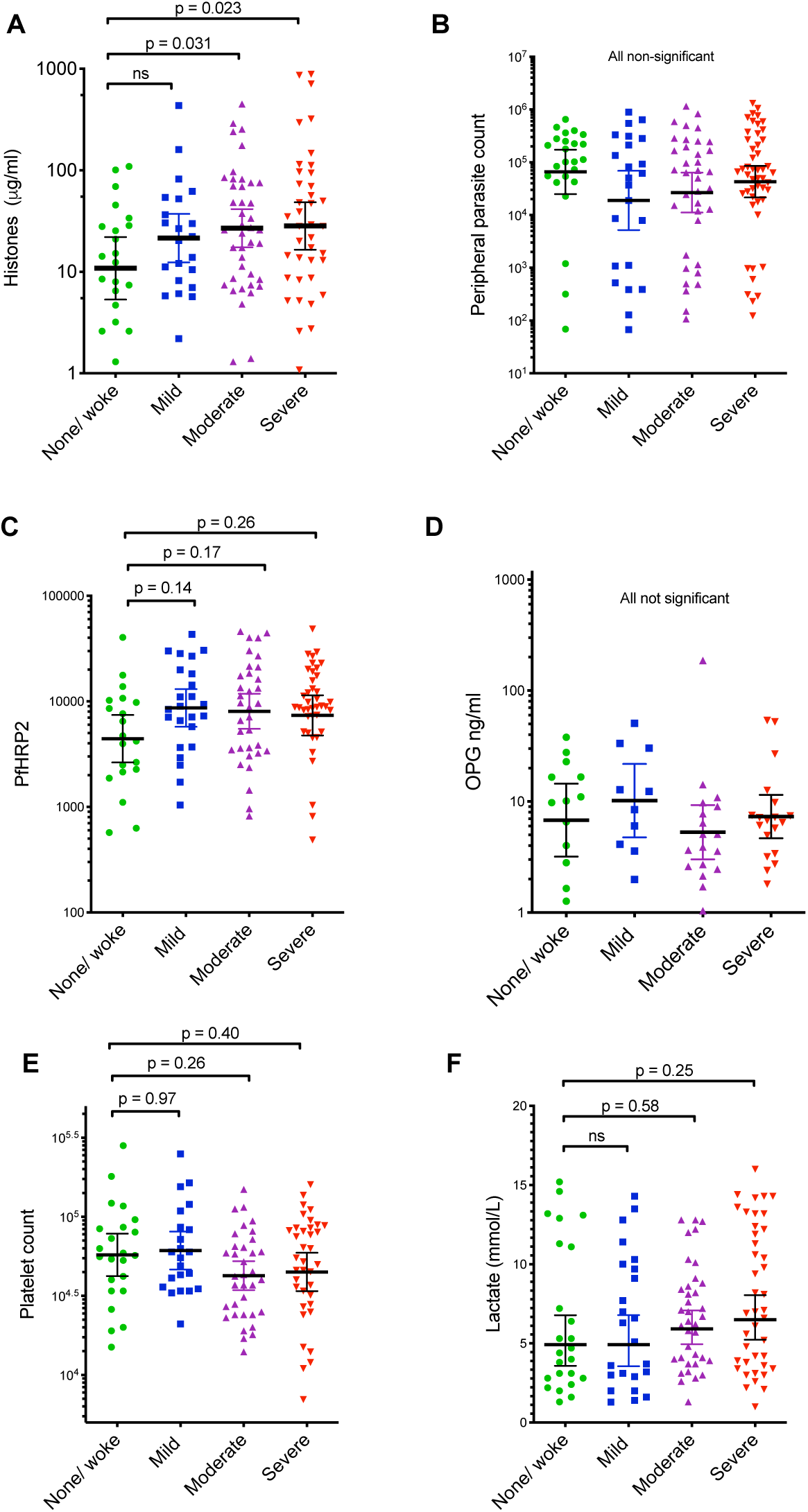
Histones but not other laboratory factors are associated with the degree of brain swelling in CM-pos patients. PfHRP2 = *P. falciparum* histidine rich protein 2; OPG = osteoprotegrin

**Fig S2.**
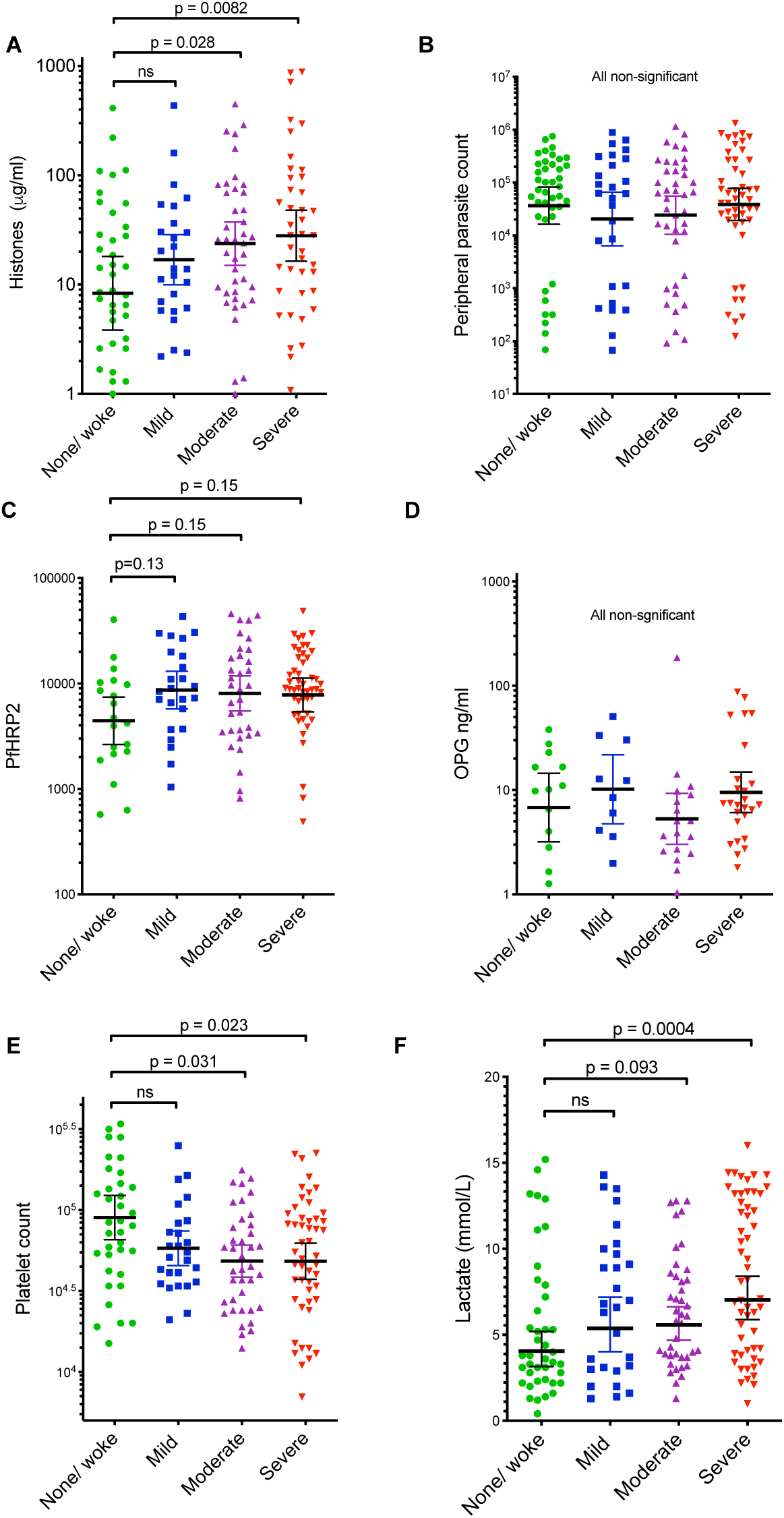
When both CM-pos and CM-neg cases are included, histones platelet count and lactate are associated with the degree of brain swelling.

**Fig. S3.**
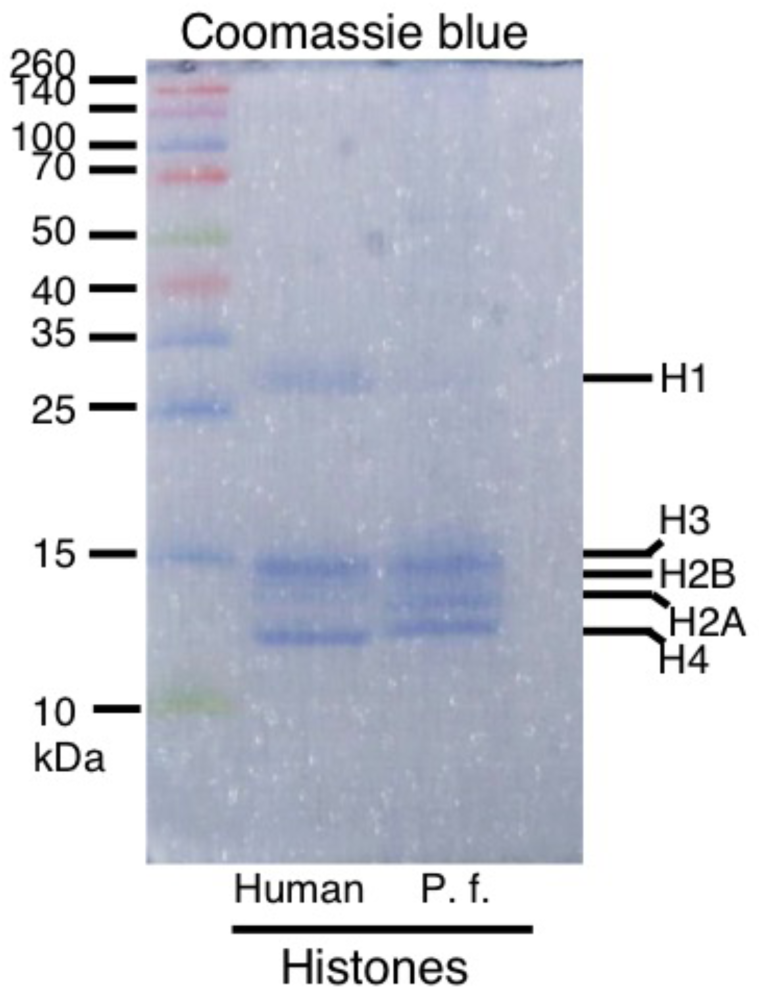
Gel showing purified Plasmodium falciparum (P. f.) and human histones. Different core histones (H2A, H2B, H3, H4) are identified by size.

**Fig. S4.**
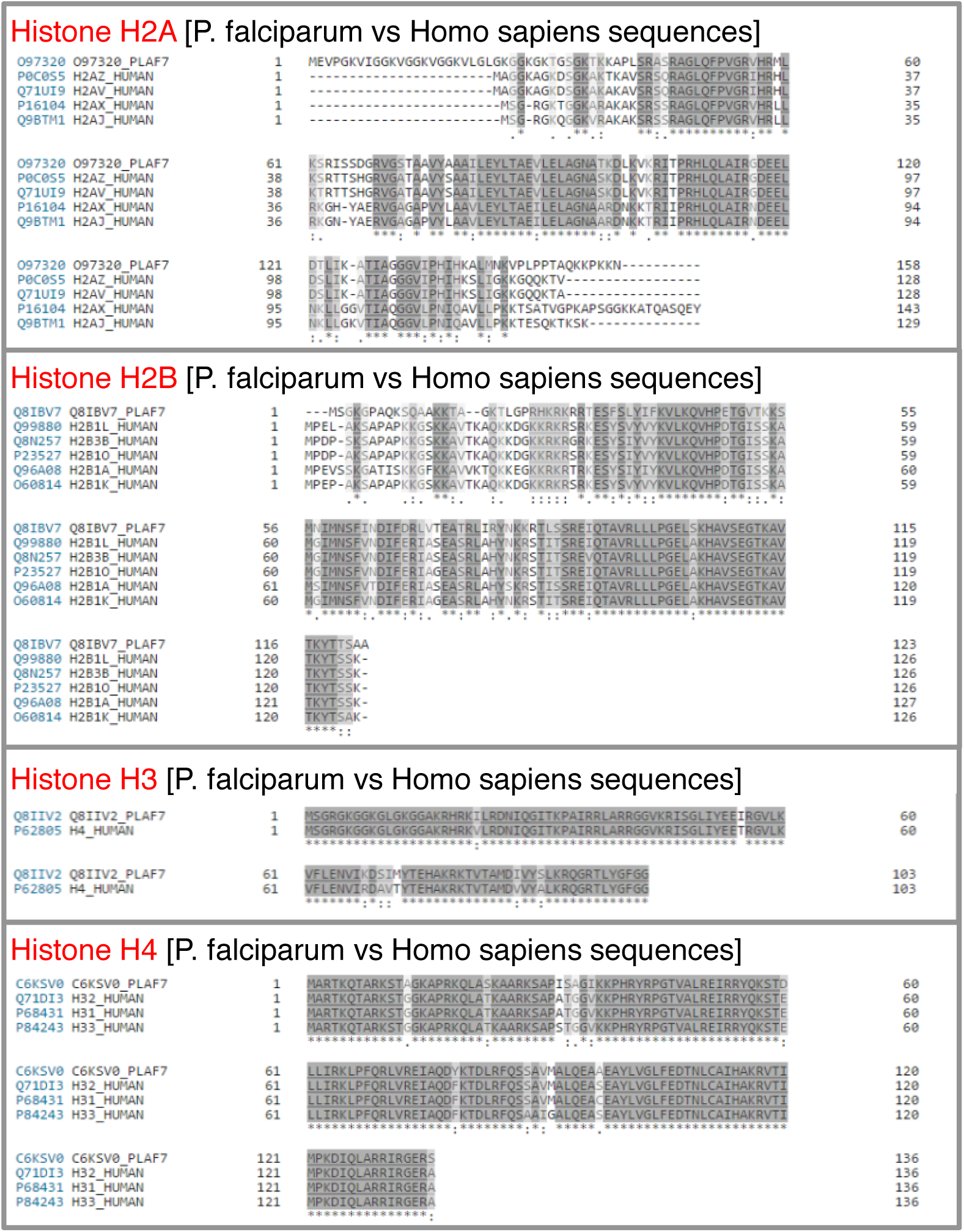
Alignment of Homo sapiens and P. falciparum histones. Amino acid sequences of individual histone variant proteins (H2A, H2B, H3 and H4) were compared between *Homo sapiens* and *P. falciparum*. Using these data, we were able to identify heterologous (species-specific) histone peptide sequences (including protein ID numbers) for further downstream analysis. Dark grey = homologous amino acids; light grey and clear = heterologous amino acids.

**Fig S5.**
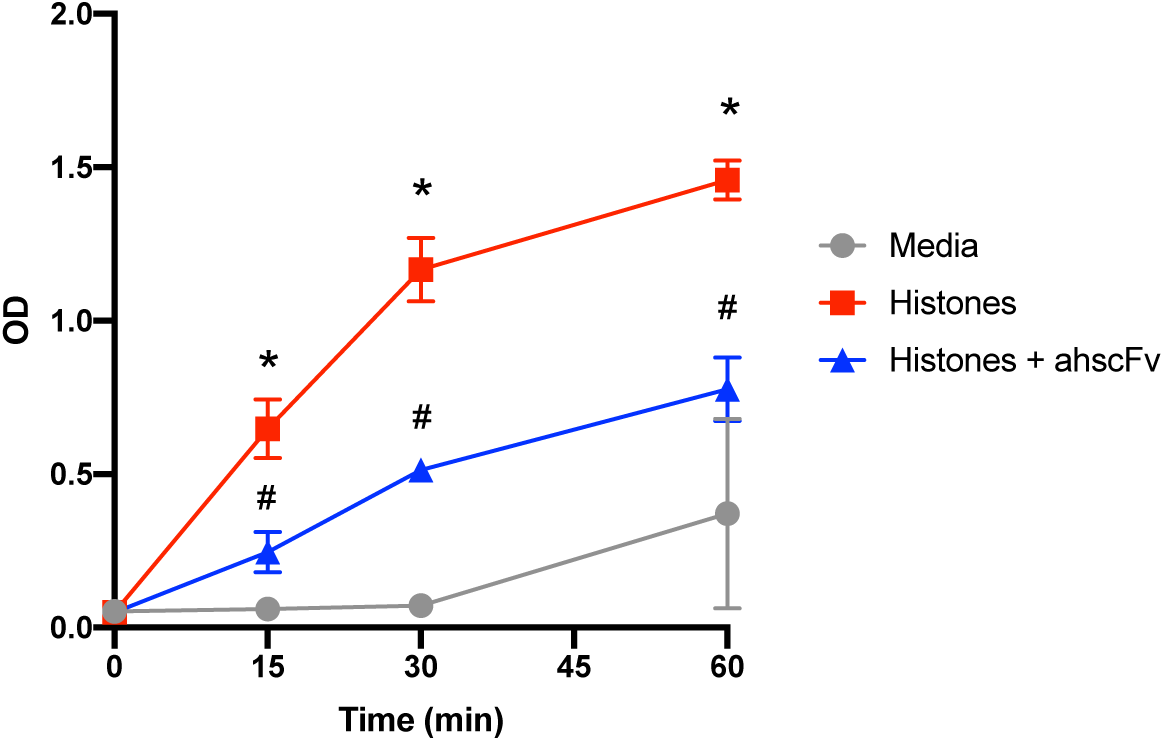
Time-course of barrier disruption of Primary human brain microvascular endothelial cells (HBMEC) by P. falciparum histones in a dual chamber system. Histone concentration 100µg/ml; Antibody concentration 200µg/ml. * = significant difference from media alone; # = significant difference from histone alone (i.e. significant protection by ahscFv).

**Table S1.**
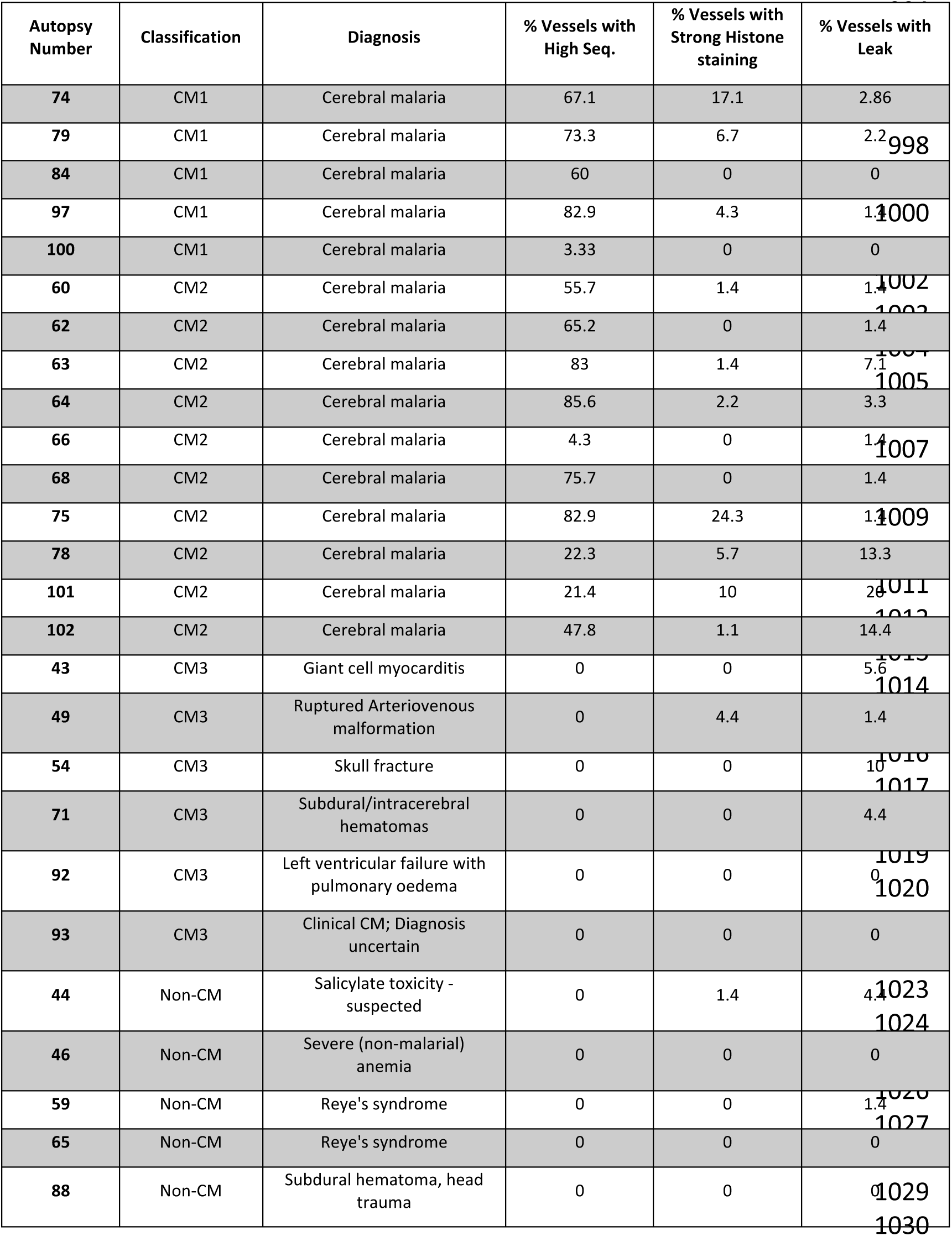
Summary of post-mortem cases. Clinical pathologist’s diagnosis at autopsy and proportion of vessels with each of: (1) high sequestration (seq; sequestration involving >50% of vessel lumen); (2) strong histone staining and; (3) leak (fibrinogen staining adjacent to a vessel).

